# Interdependent and separable functions of *C. elegans* MRN-C complex members couple formation and repair of meiotic DSBs

**DOI:** 10.1101/214015

**Authors:** Chloe Girard, Baptiste Roelens, Karl A. Zawadzki, Anne M. Villeneuve

## Abstract

Faithful inheritance of genetic information through sexual reproduction relies on the formation of crossovers between homologous chromosomes during meiosis, which in turn relies on the formation and repair of numerous double-strand DNA breaks (DSBs). As DSBs pose a potential threat to the genome, mechanisms that ensure timely and error-free DSB repair are crucial for successful meiosis. Here we identify NBS-1, the *Caenorhabditis elegans* ortholog of the NBS1 subunit of the conserved MRE11-RAD50-NBS1/Xrs2 (MRN) complex, as a key mediator of DSB repair via homologous recombination (HR) during meiosis. Loss of *nbs-1* leads to: severely reduced loading of recombinase RAD-51, ssDNA binding protein RPA and pro-crossover factor COSA-1 during meiotic prophase progression; aggregated and fragmented chromosomes at the end of meiotic prophase; and 100% progeny lethality. These phenotypes reflect a role for NBS-1 in processing of meiotic DSBs for HR that is shared with its interacting partners MRE-11-RAD-50 and COM-1 (ortholog of Com1/Sae2/CtIP). Unexpectedly, in contrast to MRE-11 and RAD-50, NBS-1 is not required for meiotic DSB formation. Meiotic defects of the *nbs-1* mutant are partially suppressed by abrogation of the non-homologous end-joining (NHEJ) pathway, indicating a role for NBS-1 in antagonizing NHEJ during meiosis. Our data further reveal that NBS-1 and COM-1 play distinct roles in promoting HR and antagonizing NHEJ. We propose a model in which different components of the MRN-C complex work together to couple meiotic DSB formation with efficient and timely engagement of HR, thereby ensuring crossover formation and restoration of genome integrity prior to the meiotic divisions.

**Significance Statement:** Double-strand breaks (DSBs) are deleterious DNA lesions, and impairment of the DSB repair machinery can lead to devastating diseases such as the Nijmegen Breakage Syndrome (NBS). During meiosis, DSBs represent a "necessary evil": they are required to promote formation of crossovers between homologous chromosomes. Crossovers in turn ensure correct chromosome inheritance during gamete formation, which is essential for viability and normal development of embryos. During meiosis, numerous DSBs are actively created, so meiotic cells must ensure that all breaks are properly repaired to ensure crossover formation and restore genomic integrity. Here we identify *C. elegans* NBS-1 as essential to properly process meiotic DSBs, both to promote crossover formation and antagonize an error-prone DSB repair pathway, thereby ensuring faithful chromosome inheritance.

## Introduction

Maintenance of genome integrity throughout cell divisions and generations is of paramount importance for organismal survival and faithful inheritance of genetic information, and multiple mechanisms have evolved to detect and repair DNA damage. Double-strand breaks (DSBs), where both DNA strands are severed, are among the most dangerous DNA lesions, as inaccurate repair of DSBs can result in genomic rearrangements, cell death and/or carcinogenesis. DSBs can be provoked by environmental sources such as radiation or chemical exposure, or can result from intrinsic cellular sources such as DNA replication errors (1).

While DSBs constitute a dangerous form of DNA damage in most cellular contexts, DSBs are deliberately induced during meiosis to promote formation of crossovers (COs) (2). Meiotic crossovers are critical for the balanced segregation of homologous chromosomes at meiosis I, as CO recombination events between the DNA molecules of homologous chromosomes, together with sister chromatid cohesion, establish physical connections between homologs (chiasmata), which in turn ensure their correct orientation toward opposite poles of the meiosis I spindle. Thus, the requirement for COs to ensure homolog segregation poses a challenge for sexually reproducing organisms, as meiotic recombination is initiated by the formation of DNA lesions that constitute a danger to genomic integrity. By the end of meiosis, all DSBs must be accurately repaired to (i) ensure CO formation and proper chromosome segregation, and (ii) guarantee that genome integrity is restored prior to cell division.

Meiotic DSBs are specifically induced by the conserved topoisomerase VI-like protein SPO11 (3–5). The SPO11 protein remains covalently bound to both broken DNA ends after the break occurs, and has to be removed through a process called resection for the DSB to be repaired. Resection is initiated by an endonucleolytic cleavage that leads to the release of SPO11 attached to a small oligonucleotide (6) and results in a short 3' single stranded DNA (ssDNA) tail. Further resection of the 5' end produces a longer ssDNA tail (7), which recruits DNA strand exchange proteins DMC1 and/or RAD51 to stimulate invasion of an homologous DNA duplex and repair of the DSB by homologous recombination (HR) (8). The first DNA cleavage event is dependent on endonuclease activity of the conserved MRN/X complex composed of MRE11, RAD50 and NBS1/XRS2, as well as on the COM1/Sae2/CtIP/Ctp1 protein, which associates with MRN/X (9). Analysis of budding yeast meiosis shows that in the second step, the Exo1 exonuclease joins in to extend the resected tracts and produce the long 3'-ssDNA-tailed intermediates (10).

An alternative mechanism for DSB repair (DSBR) is the non-homologous end-joining (NHEJ) pathway, which involves protection of the broken ends by the Ku70/Ku80 heterodimer ring (11). Binding of Ku prepares DSBs for direct ligation between broken DNA ends with little or no homology, an inherently error-prone process (12). In cases where multiple DSBs on different chromosomes are present in the same cell, as occurs during meiosis, end-joining can result in chromosome translocations. In contrast to NHEJ, homologous recombination is generally considered an error-free pathway of DSBR as it uses a homologous DNA template to repair the broken molecule. A strong body of evidence indicates that there is competition between the HR and NHEJ pathways for repair of DSBs, raising the question as to how pathway choice is regulated (12). Initiation of resection by the MRN/X complex and Com1/Sae2/CtIP/Ctp1 appears to be critical for this decision, as it commits cells to homology-dependent repair (7). Interestingly, evidence from *C. elegans* indicates that such competition occurs even during meiosis, where it is absolutely critical for DSB repair to occur exclusively by HR (13, 14). Thus, efficient coupling of DSB formation and DSB resection is of paramount importance for ensuring a successful outcome of meiosis.

MRE11 and RAD50 are highly conserved in eukaryotes. MRE11 is the nuclease subunit of the complex, while RAD50, which belongs to the Structural Maintenance of Chromosomes (SMC) family of proteins, is required for regulating MRE11 nuclease activity in an ATP-dependent manner and may also be important for tethering of DNA ends (7). Nbs1/Xrs2 is the least conserved member of the MRN/X complex, and the high sequence divergence between mammalian NBS1 and yeast XRS2 had precluded the identification of orthologs in many species (15), including *C. elegans*.

Here, we report the identification of the previously elusive *C. elegans* NBS-1 ortholog based on a role in meiotic recombination revealed by a mutant screen. Unexpectedly, we found that the requirements for NBS-1 during meiosis are distinct from those of its complex partners. In contrast to MRE-11 and RAD-50, which are required both for formation and resection of meiotic DSBs (14, 16, 17), NBS-1 is required for DSB resection but is dispensable for DSB formation. We further found that NBS-1 (like MRE-11 (14)) is not only important for promoting resection and HR but also for antagonizing NHEJ during meiosis. This latter characteristic is shared with COM-1 (13, 18), a partner of the MRN complex, but our data reveal distinct roles for NBS-1 and COM-1 in promoting HR and antagonizing NHEJ. Our results support a model in which different components of the MRN-C complex work together during meiosis to couple formation and repair of meiotic DSBs to both (i) promote efficient and timely DSB resection to promote HR and (ii) antagonize NHEJ to ensure genome stability.

## Results

### Identification of the C. elegans NBS-1 ortholog

We isolated the initial *nbs-1(me102)* mutant allele in a genetic screen for

*C. elegans* mutants with altered numbers of GFP::COSA-1 foci, which mark the sites of COs in *C. elegans* germ cells at the late pachytene stage of meiotic prophase (Figure 1A). As each chromosome pair normally undergoes only a single CO during *C. elegans* meiosis, wild-type late pachytene nuclei consistently exhibit 6 GFP::COSA-1 foci, one for each pair of homologs (19). Further, DAPI staining of WT oocytes at diakinesis, the last stage of meiotic prophase I, reveals 6 well-resolved DAPI bodies corresponding to the 6 pairs of homologs linked by chiasmata (6 bivalents). The *nbs-1(me102)* mutant was isolated based on observation of a severe reduction in the number of GFP::COSA-1 foci by live imaging (Figure 1A), indicating impairment of meiotic recombination. DAPI staining of diakinesis oocytes in the *nbs-1(me102)* mutant further revealed frayed, aggregated and/or fragmented chromosomes (Figure 1B & 1C), indicative of defects in DNA repair. Moreover, *me102* homozygous hermaphrodites produced no viable progeny (0 survivors/1575 eggs laid, Table 1).

**Table 1:** Quantitation of progeny viability and male frequency

| Genotype | Average number of eggs laid ± SE<br>(number of broods) | % viable adults<br>(total number of eggs laid) | % males |
| --- | --- | --- | --- |
| Wild type | 287 ± 5 (n=10) | 100% (2878) | 0.05% |
| *nbs-1* | 157 ± 8 (n=10) | 0% (1575) | NA |
| *spo-11* | 69.4 ± 9 (n=9) | 9% (516) | 27% |
| *nbs-1; spo-11* | 53.8 ± 9 (n=8) | 15% (366) | 40% |
| *cku-80* | 204 ± 15 (n=7) | 100% (1427) | 0.1% |
| *exo-1* | 225 ± 17 (n=10) | 100% (2251) | 0.7% |
| *cku-80 exo-1* | 206 ± 19 (n=10) | 100% (2065) | 0.004% |
| *nbs-1 ; cku-80* | 158 ± 22 (n=19) | 3.1% (3007) | 14% |
| *nbs-1; exo-1* | 4 ± 6 (n=19) | 0% (78) | NA |
| *nbs-1; cku-80 exo-1* | 45.5 ± 7 (n=10) | 0% (455) | NA |

**Figure 1:**
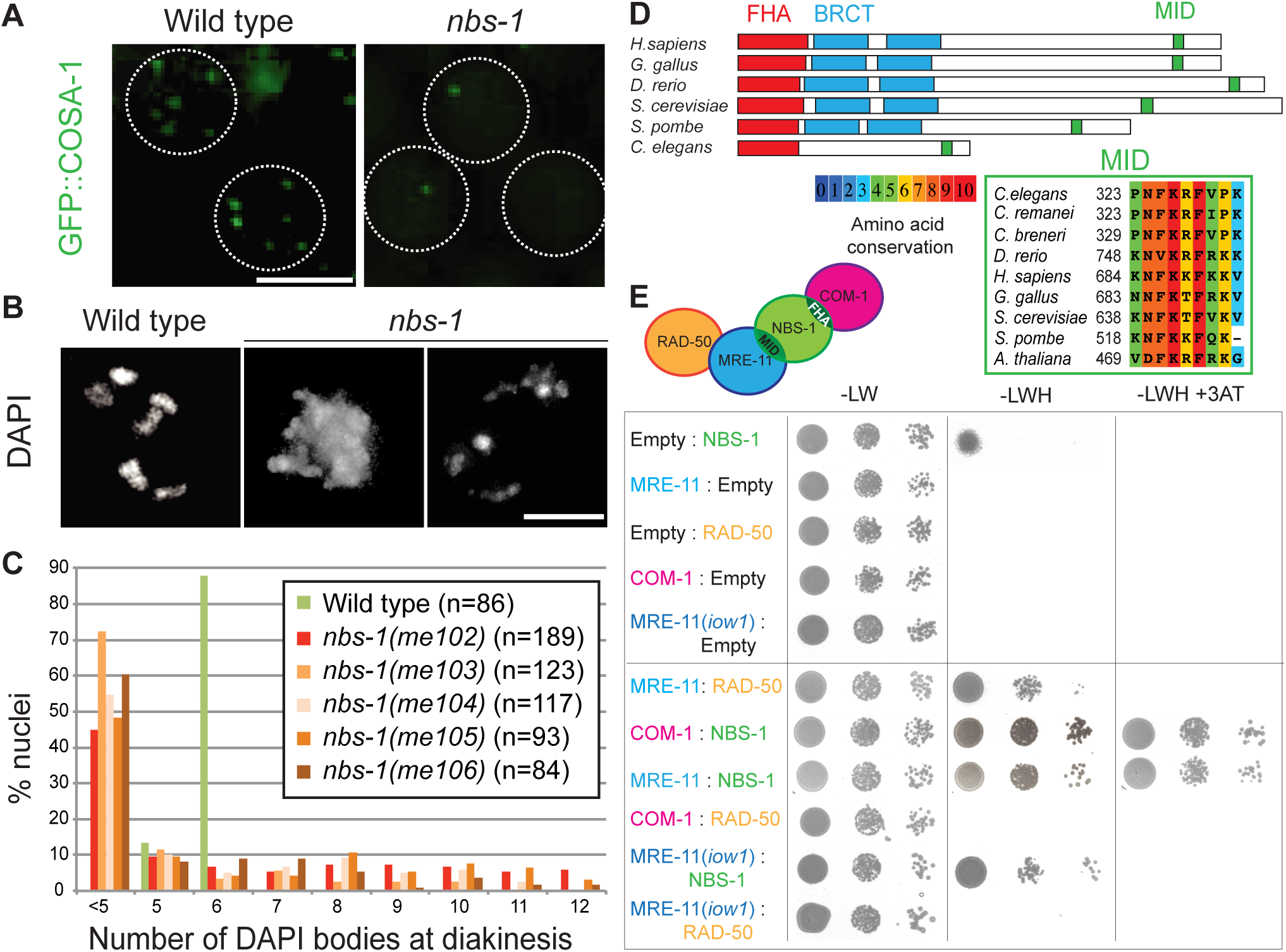
Identification of the *C. elegans nbs-1* ortholog, based on its requirement for meiotic DNA repair. **A)** GFP::COSA-1 foci in late pachytene nuclei of live worms. Each wild-type nucleus has 6 foci (corresponding to the 6 CO sites), while the *nbs-1* nuclei usually have one or zero foci. Scale bar: 5μm. **B)** Images of DAPI stained chromosomes from individual diakinesis-stage oocytes. The wild-type oocyte displays six DAPI bodies corresponding to the six pairs of homologs connected by chiasmata, while the *nbs-1(me102)* mutant oocytes display chromosome aggregates (less than 5 DAPI bodies), indicative of defective DNA repair. Scale bar: 5μm**. C)** Graphs showing frequencies of diakinesis-stage oocytes with the indicated number of DAPI bodies in wild type worms and worms homozygous for *nbs-1* mutant alleles (see also Figure S1). **D)** Top: Schematic depicting the *C. elegans* NBS-1 protein and its orthologs in other species. NBS-1 contains the conserved Forkhead Associated (FHA) domain and the MRE-11 Interacting Domain (MID) but lacks the tandem BRCT domains. Bottom: Alignment showing conservation of the MID among members of the NBS1/Nibrin protein family. **E)** Yeast two-hybrid assay revealing interactions between NBS-1 and its cognate partners. Interaction between prey proteins fused with the GAL4 activation domain (left) and the baits fused with the LexA DNA binding domain (right) assayed by growth on media lacking histidine (-LWH); growth in the presence of 3-AT, a competitive inhibitor of His3p, indicates strong interaction. Serial dilutions are spotted (1, 1:100; 1:1000).

The causal mutation was mapped to a ~6.8cM region on chromosome *II*. Whole genome sequencing of a 3X backcrossed strain identified two mutations in the interval, one being a nonsense mutation in the *C09H10.10* gene (see Materials and Methods). Insertion/deletion mutant alleles were generated using CRISPR technology, creating early frame-shifts and stop codons in *C09H10.10* (Figure S1). All four CRISPR-derived alleles recapitulated the diakinesis and progeny inviability phenotypes of *me102*, confirming that disruption of *C09H10.10* is responsible for the observed phenotypes (Figure 1C) and suggesting that all 5 mutant alleles (*me102-6)* of *C09H10.10* are likely null alleles.

The predicted C09H10.10 protein contains a conserved FHA domain (<u>F</u>ork<u>h</u>ead-<u>a</u>ssociated domain, Figure S1A) at the N-terminus, and PSI-BLAST searches initiated using C09H10.10 as the query sequence detected homology with the *Danio rerio* Nibrin protein, a predicted ortholog of mammalian NBS1. *Caenorhabditis* C09H10.10 orthologs lack the tandem BRCT domains found adjacent to the FHA domain in previously-recognized NBS1/Xrs2 orthologs (15). However, a small but highly conserved MRE11 interacting domain (MID) discovered in *S. pombe* Nbs1 (20) is clearly recognizable near the C-terminus of C09H10.10 (Figure 1D). These features, coupled with functional data presented below, identify C09H10.10 as the *C. elegans* NBS1 ortholog, hereafter referred to as NBS-1.

Yeast two-hybrid assays revealed interactions between *C. elegans* NBS-1 and MRE-11 and between NBS-1 and COM-1, and confirmed the previously-reported interaction between MRE-11 and RAD-50 (21), recapitulating the interaction network described in other species (Figure 1E) (22). Homozygous *nbs-1* worms from heterozygous parents are fully viable and do not show any developmental phenotype in normal growth conditions, which allowed us to investigate the role of NBS-1 in DSB repair during meiosis.

### C. elegans NBS-1 is required for DSB repair but not for DSB formation

Multiple lines of evidence indicate that the presence of chromosome aggregates in *nbs-1* mutants reflects a defect in repair of the SPO-11-dependent DSBs that serve as the initiating events of meiotic recombination. The *spo-11* mutant lacks meiotic DSBs, resulting in lack of COs and chiasmata, which is reflected by the presence of 12 unattached chromosomes (univalents) at diakinesis (23). In contrast to the *nbs-1* single mutant, the *nbs-1; spo-11* double mutant displayed the canonical *spo-11* phenotype, exhibiting 12 DAPI bodies at diakinesis (Figure 2A & B) and producing a few percent viable progeny due to occasional euploid embryos arising from erratic segregation of intact chromosomes at meiosis I (Table 1). This indicates that the complete progeny lethality and the aggregated/fragmented chromosomes in diakinesis nuclei observed in *nbs-1* mutants are a consequence of meiotic DSBs. Further, while introduction of exogenous DSBs rescued the chiasma formation defect of the *spo-11* single mutant, as shown by diakinesis nuclei displaying 6 DAPI bodies (23), frayed and aggregated chromosomes were observed following irradiation in *nbs-1; spo-11* diakinesis nuclei, demonstrating impaired repair of DSBs, whether SPO-11-dependent or exogenously induced, in absence of NBS-1 (Figure 2A & 2C).

**Figure 2:**
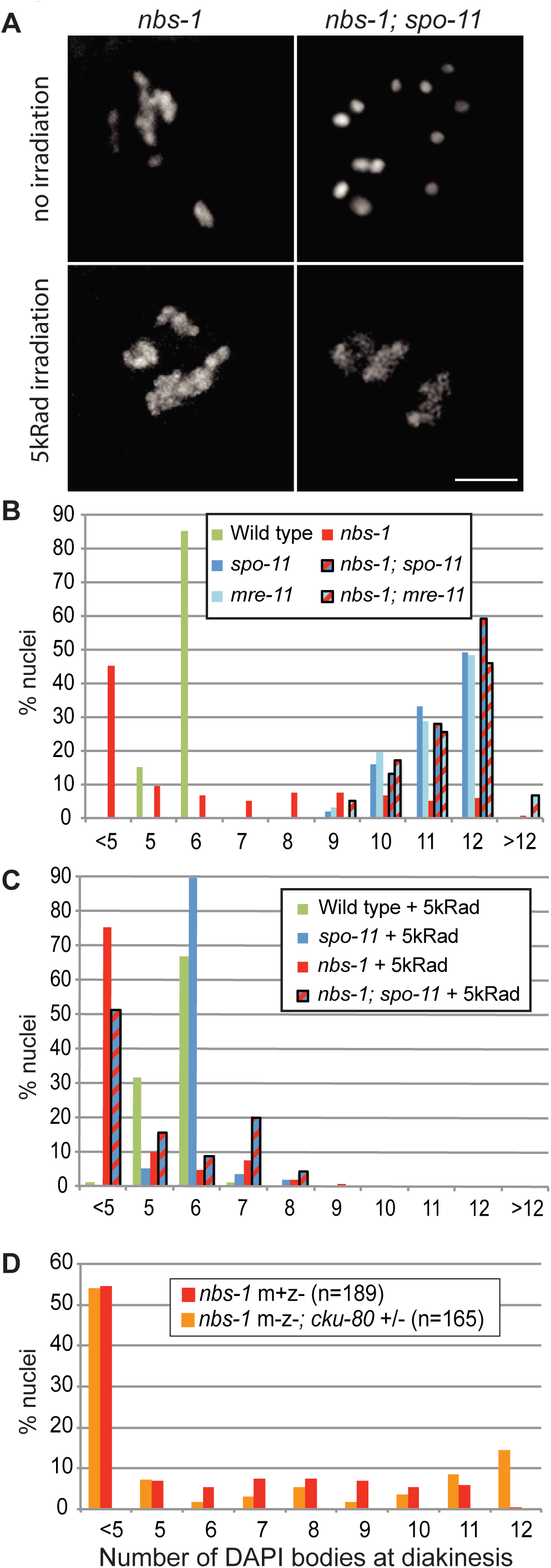
*C. elegans* NBS-1 is required for meiotic DSB repair but dispensable for DSB formation. **A)** DAPI stained diakinesis oocytes from *nbs-1* and *nbs-1; spo-11* mutants worms unirradiated or exposed to 5kRad γ-irradiation. In contrast to the abnormal chromosome aggregates present in the *nbs-1* oocytes (with or without irradiation), 12 intact chromosomes (univalents) are observed in the unirradiated *nbs-1; spo-11* oocyte. Aggregation of chromosomes is however observed in the *nbs-1; spo-11* oocyte upon introduction of exogenous DSBs by irradiation. Scale bar: 5μm. **B)** Quantification of the number of DAPI bodies in diakinesis nuclei. Fewer than 5 countable DAPI bodies reflects aggregation of chromosomes, whereas 12 DAPI bodies typically reflects intact univalents. Numbers of nuclei counted: wild type n=92, *spo-11* n=37, *nbs-1(me102)* n=42, *nbs-1; spo11* n=39, *mre-11* n=66, *nbs-1; mre-11* n=59. **C)** Quantification of DAPI bodies as in (B) following exposure of worms to 5kRad γ-irradiation, showing that irradiation-induced breaks rescue chiasma formation in the *spo-11* mutant but induce chromosome aggregation in *nbs-1; spo-11* mutant oocytes. Numbers of oocytes counted: wild type n=105, *spo-11* n=57, *nbs-1(me102)* n=144, *nbs-1; spo-11* n=45. **D)** Graph showing indistinguishable profiles of diakinesis DAPI body counts in *nbs-1(me102)* mutant worms derived from heterozygous *nbs-1/+* mothers (m+z-) and *nbs-1* m-z-mutant worms, which were derived from a cross using homozygous *nbs-1; cku-80* double mutant mothers (m-z-).; see Figure S2 for more details.

Our finding that meiotic DSBs are still formed in *nbs-1* null mutant worms was unexpected, as previous studies had shown that the two other partners of the MRN complex, MRE-11 and RAD-50, are required for both DSB formation and DSB repair during *C. elegans* meiosis (14, 16, 17). Further, our own data showing that the *nbs-1; mre-11* double mutant displays 12 intact univalents (and no aggregates) at diakinesis indicates that MRE-11 is still required for meiotic DSB formation in an *nbs-1* mutant background (Figure 2B). Since our analyses were conducted using *nbs-1/nbs-1* worms derived from *nbs-1/+* mothers (m+z-animals), we considered the possibility that DSB formation in the germ lines of *nbs-1* m+z-animals could be the consequence of residual maternal NBS-1 protein. Although *nbs-1/nbs-1* mutant progeny from *nbs-1/nbs-1* mothers (m-z-) are normally completely inviable, we devised a crossing strategy that enabled us to generate some viable *nbs-1/nbs-1* m-z-worms (Figure S2 and see below); these m-z-*nbs-1* worms displayed the same phenotype of aggregated chromosomes at diakinesis as their m+z-counterparts, indicating proficiency for DSB formation but deficiency in DSB repair (Figure 2D). These results show that *C. elegans* NBS-1, unlike MRE-11 and RAD-50, is dispensable for DSB formation, and that MRE-11 and RAD-50 can act independently of NBS-1 to promote meiotic DSB formation.

These results are reminiscent of a previously described separation-of-function mutant *mre-11(iow1)* that is proficient for DSB formation but not DSB repair (14). We thus tested whether the *iow1* mutation might perturb the interaction between MRE-11 and NBS-1, impairing DSB repair, while leaving the interaction between MRE-11 and RAD-50 intact to enable DSB formation. However, yeast two-hybrid assays showed that the *mre-11(iow1)* mutation weakened but did not eliminate the interaction between MRE-11 and NBS-1, and disrupted the interaction between MRE-11 and RAD-50 (Figure 1E). This result suggests that the interaction interface between MRE-11 and RAD-50 might not be as crucial for DSB formation as it is for DSB repair.

### NBS-1 is essential for DSB resection and loading of RAD-51 and RPA-1

Normal repair of meiotic DSBs requires ends to be processed so that they can engage in HR-mediated repair, both to form COs and to restore genome integrity. More specifically, SPO-11 protein-DNA adducts must be removed from 5' ends through an endonucleolytic process. Furthermore, DSB ends must be further resected to yield 3' ssDNA tails that can recruit DNA strand exchange proteins such as RAD-51 to mediate invasion of a homologous DNA template.

As NBS1 is a member of the MRN complex involved in DSB resection in other species, we assessed the ability of *nbs-1* mutants to process SPO-11-dependent DSBs by: i) simultaneous visualization of RAD-51 and a tagged version of RPA-1 (RPA-1::YFP (24)), a component of eukaryotic ssDNA binding protein RPA, following nuclear spreading (Figure 3A); and ii) quantification of RAD-51 foci in whole-mount gonads representing a time course of nuclei entering and progressing through meiosis (Figure 3C). In wild-type *C. elegans* meiosis, RAD-51 foci appear during zygotene and early pachytene following DSB resection and become numerous by mid-pachytene before disappearing by late-pachytene, indicative of efficient DSB repair (25, 26). When observed using structured illumination microscopy (SIM), RAD-51 foci typically appear as doublets, reflecting resection of both DSB ends (Figure 3B; (27)). In addition, RPA-1 foci, most of which represent post-strand-exchange recombination intermediates, rise in abundance and accumulate to higher levels than RAD-51 foci before decreasing and disappearing during late pachytene (27).

**Figure 3:**
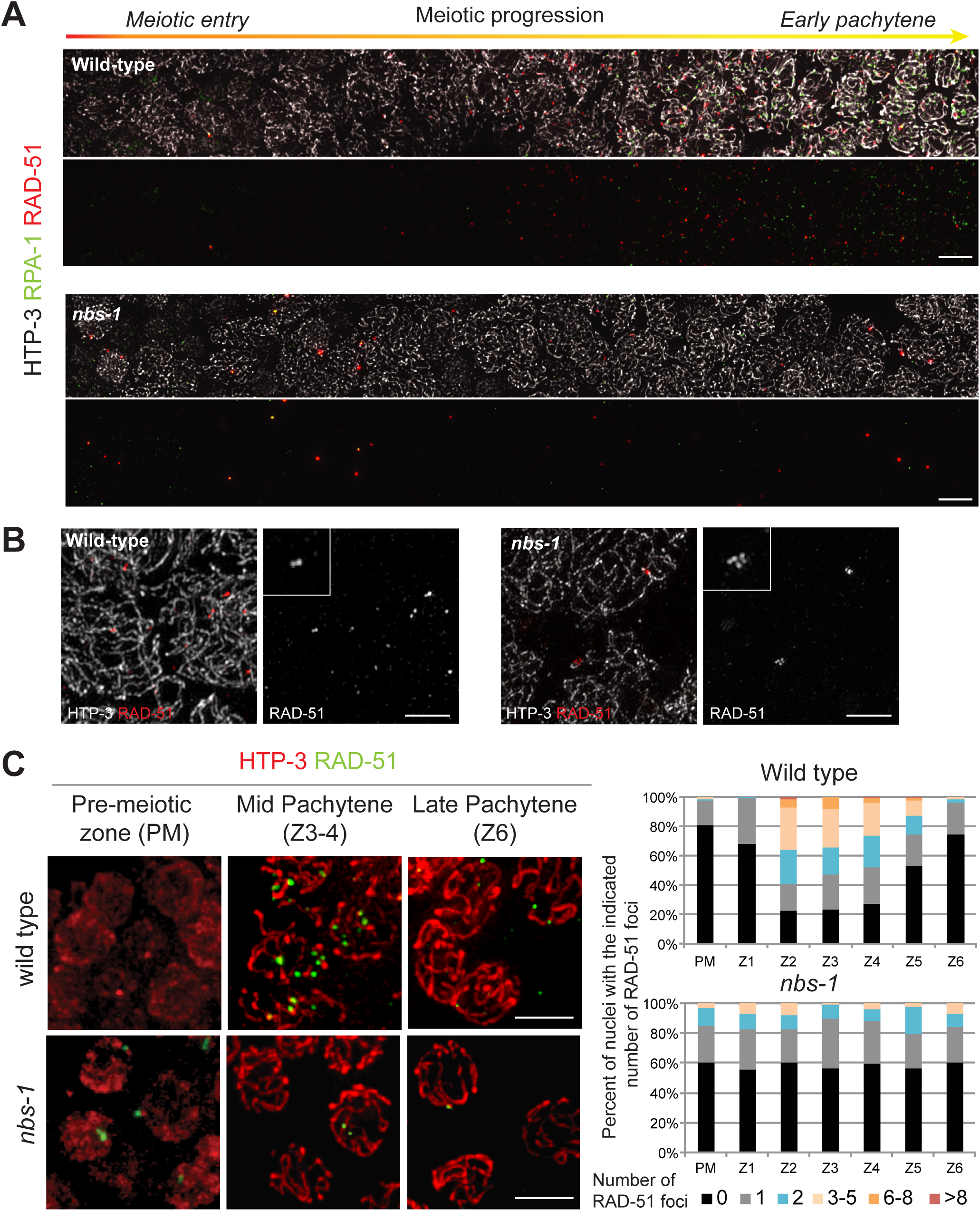
Abundance of RAD-51 and RPA foci is greatly reduced in the absence of NBS-1. **A)** Images of nuclei from spread gonads from wild-type and *nbs-1(me103)* mutant worms, immunostained for chromosome axis protein HTP-3 (greyscale), recombinase RAD-51 (red) and a YFP-tagged version of RPA-1 (green), a subunit of the ssDNA binding protein RPA. Gonad segments depicted include a few pre-meiotic nuclei (diffuse HTP-3 staining) and meiotic prophase stages (recognized by linear tracks of HTP-3) ranging from meiotic entry through early pachytene. During early meiotic prophase progression, RAD-51 and RPA-1 foci rise in abundance in the wild-type gonad but not in the *nbs-1* mutant gonad, where only a subset of nuclei distributed throughout the region have one or a few bright foci. Scale bar: 15 μm. **B)** Structured illumination microscopy (SIM) images of RAD-51 foci from the spread nuclei in (A). Scale bar: 2μm. Inset images show the doublet (or singlet) organization characteristic of RAD-51 foci at meiotic DSB sites during wild-type meiosis (left) and the more complex organization of RAD-51 foci detected in *nbs-1* nuclei (right), which are thought to reflect abnormalities arising during replication that persist into meiotic prophase. **C)** Left: Representative images of germ cell nuclei in whole-mount preparations immunostained for HTP-3 (red) and RAD-51 (green), illustrating both the higher numbers of foci detected in mid-pachytene nuclei in the wild-type and the abnormal foci detected in a subset premeiotic nuclei in the *nbs-1* mutant Scale bar: 10μm. Right: quantification of the numbers of RAD-51 foci in nuclei (from the whole-mount preparations) in seven consecutive zones along the distal-proximal axis of the gonad from the pre-meiotic region (PM) through the end of pachytene (Z6; see Figure S3A).

Consistent with previous reports indicating a role for MRE-11 and RAD-50 in the processing of meiotic DSBs (14, 16, 17), we found that the *nbs-1* mutant is impaired for RAD-51 focus formation, exhibiting an overall reduction in the abundance of RAD-51 foci and an absence of a mid-pachytene peak in foci numbers (Figure 3A & 3C). Further, the abundance of RPA-1 foci was also severely reduced in the *nbs-1* mutant. Thus, *nbs-1* mutant germ cells do not accumulate post-strand-exchange recombination intermediates (as occurs during wild-type meiosis), nor do they accumulate RPA-coated ssDNA ends (as occurs in *brc-2* mutants, which are competent for DSB resection but defective in RAD-51 loading (28)). Together these data indicate that NBS-1 is essential for meiotic DSB resection.

Whereas numbers of RAD-51 and RPA-1 foci were reduced overall, *nbs-1* mutants displayed an increased number of foci in the premeiotic zone (PM, Figure 3A and 3C), consistent with a role for MRN in repairing and/or preventing accumulation of DNA damage during DNA replication during mitosis before meiotic entry (17, 29). Supporting this interpretation, we found that an *nbs-1; spo-11* double mutant exhibited higher levels of residual RAD-51 foci (0.44 ± 0.75 foci per nucleus in zones 1 through 6, n=727) than the *spo-11* single mutant (0.21 ± 0.58, n=1094; Mann-Whitney p<10^-4^) (Figure S3), suggesting that many of the residual RAD-51 foci detected in *nbs-1* meiotic nuclei reflect DNA damage that was not of meiotic origin. Further, SIM imaging revealed that RAD-51 foci in the *nbs-1* mutant exhibit abnormal structure (Figure 3B). In contrast to the doublet or singlet foci observed in wild-type germ cells (27), RAD-51 foci in the *nbs-1* mutant are typically larger and more complex, both in the premeiotic zone and throughout meiotic prophase, consistent with abnormalities arising during mitotic cell cycles or meiotic DNA replication and persisting following meiotic prophase entry. However, we also found that the residual level of RAD-51 foci in the *nbs-1* single mutant (0.63 ± 0.88, n=618 nuclei) was higher than in the *nbs-1; spo-11* double mutant (Mann-Whitney p<10^-4^) (Figure 3C & S3); this suggests that although meiotic DSB resection is strongly impaired, some SPO-11-generated breaks may nevertheless load RAD-51 in the absence of NBS-1.

### NBS-1 functions both to counteract non-homologous end joining and to promote efficient HR

DNA repair pathway choice is crucial for cellular and organismal survival: non-homologous end joining (NHEJ) and HR have been shown to occur cooperatively, competitively or as backup mechanisms for DSB repair in various contexts (12). As previous reports had implicated MRE-11 and COM-1 in antagonizing NHEJ (13, 14), we tested the hypothesis that the meiotic defects observed in the *nbs-1* mutant might reflect inappropriate use of NHEJ for the repair of meiotic DSBs.

We found that mutation of *cku-80*, which encodes the worm ortholog of KU80 essential for NHEJ, partially alleviated multiple *nbs-1* defects (Figure 4). In contrast to the aggregated chromosomes present in the *nbs-1* single mutant, diakinesis chromosomes more frequently appeared as individual univalents or bivalents in the *nbs-1; cku-80* double mutant (Figure 4A). This partial restoration of chromosome integrity was accompanied by a partial restoration of GFP::COSA-1 foci in late pachytene (Figure 4B). While the *nbs-1* mutant displayed an average of 1.4 ± 1.3 (n=150) foci per nucleus, the *nbs-1; cku-80* double mutant averaged 4.7 ± 1.5 GFP::COSA-1 foci per nucleus (n= 133, Mann-Whitney p<10^-4^). We also observed a partial rescue of progeny viability, with an average of 3.1% progeny survivorship from *nbs-1; cku-80* animals (93/3007 eggs laid) compared to 0% from *nbs-1* animals (0/1575, Table 1). The substantial rescue of progeny viability, chromosome integrity and GFP::COSA-1 focus formation together indicate a role for NBS-1 in preventing inappropriate utilization of NHEJ during meiosis.

**Figure 4:**
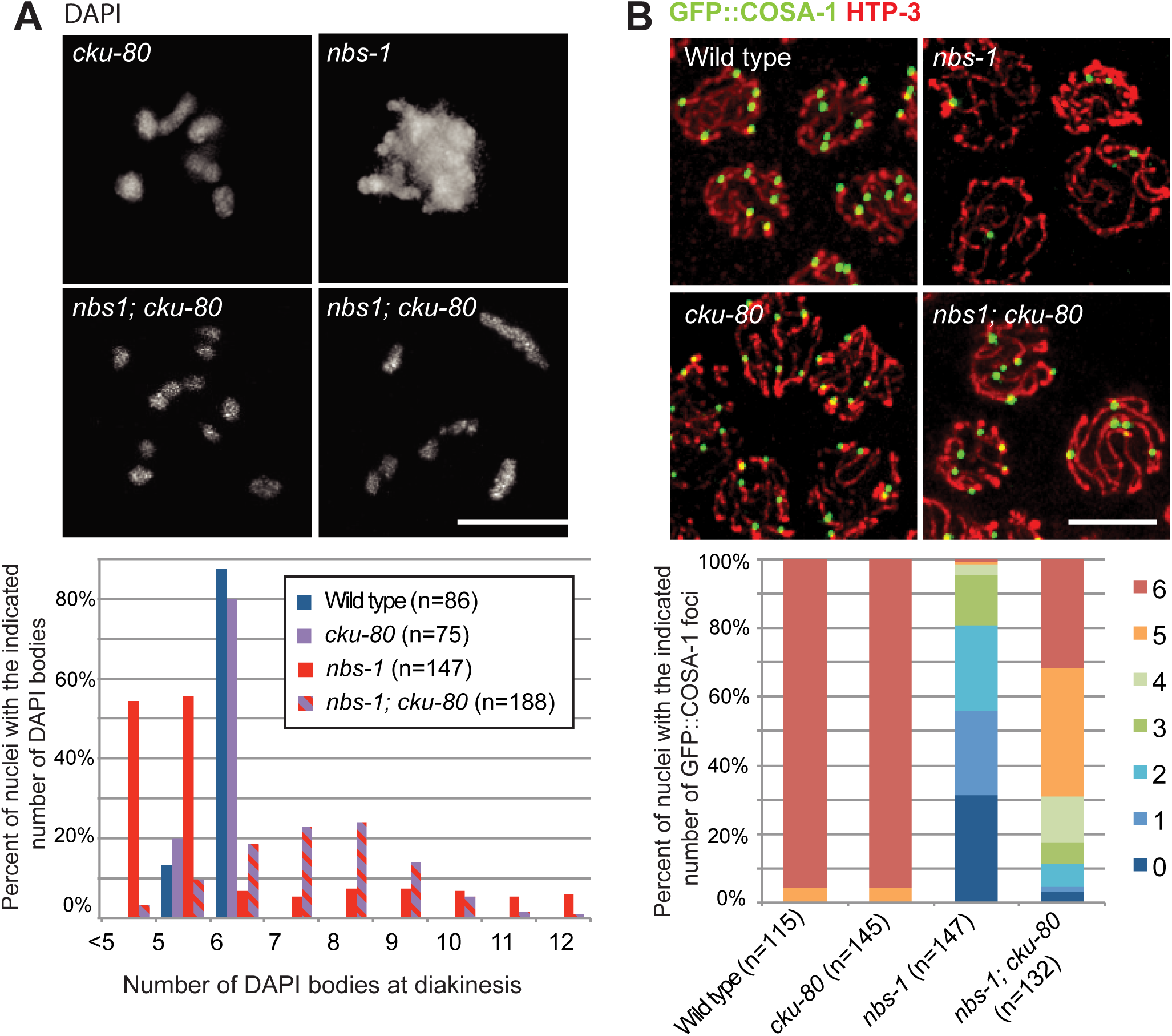
NBS-1 antagonizes NHEJ and promotes efficient HR. **A)** Top: Images of individual DAPI stained diakinesis oocyte nuclei showing that the chromosome aggregation phenotype of the *nbs-1* single mutant is suppressed in *nbs-1; cku-80* double mutant, which instead displays a mixture of bivalents and univalents. Scale bar: 5µm. Bottom: quantification of DAPI bodies in diakinesis nuclei. **B)** Top: Immunolocalization of HTP-3 (red) and GFP::COSA-1 (green) in late pachytene nuclei in whole-mount preparations (scale bar 10µm). Bottom: Stacked bar graph showing the percentages of nuclei with the indicated numbers of COSA-1 foci, showing that COSA-1 focus formation is partially restored in the *nbs-1; cku-80* double mutant compared to the *nbs-1* single mutant.

Although inactivation of *cku-80* attenuated the meiotic defects of the *nbs-1* mutant, the rescue was not complete. This result could reflect either (i) an additional role for NBS-1 in promoting efficient HR beyond antagonizing NHEJ; or (ii) a deficit of DSBs compared to wild type, which could yield a deficit in CO number. We ruled out the latter hypothesis by exposing *nbs-1; cku-80* worms to 5kRad γ-irradiation to introduce an excess of DSBs. While this dose is more than sufficient to restore chiasmata in the *spo-11* mutant background (Figure 2C and (19, 23), it did not improve chiasma formation in the *nbs-1; cku-80* mutant (Figure S4). This result indicates that DSBs are not limiting for CO formation in the *nbs-1; cku-80* mutant, and instead implies that recombination intermediates cannot be efficiently processed into COs in absence of NBS-1, even when CKU-80 is absent.

### NBS-1 is required for a timely resection of DSBs to engage HR

Examination of the timing of appearance of RAD-51 and RPA-1 foci in *nbs-1; cku-80* double mutant indicated a role for NBS-1 in promoting timely resection of DSBs, even in absence of NHEJ (Figure 5). The *nbs-1; cku-80* double mutant differed from both the *cku-80* single mutant, which exhibits wild-type dynamics of RAD-51 and RPA-1 foci with an enrichment in mid-pachytene (Figure 5A), and from the *nbs-1* single mutant, which displays low levels of both types of foci throughout meiosis I (Figure 3C). Instead, numbers of RAD-51 and RPA-1 foci in *nbs-1; cku-80* remained low throughout most of meiotic prophase, but then rose in abundance during late pachytene (Zone 6, Figure 5C and 5D), similar to what was reported for RAD-51 foci in the *mre-11(iow1); cku-80* double mutant (14). The majority of these late RAD-51 foci appeared as doublets when resolved by SIM imaging (Figure 5B), as they do in wild type during early pachytene, consistent with the presence of resected meiotic DSB ends. These results suggest that some resection can occur in the absence of NBS-1, but only if NHEJ is abrogated. Moreover, this NBS-1 independent mode of resection appears largely restricted to late pachytene and early diplotene.

**Figure 5:**
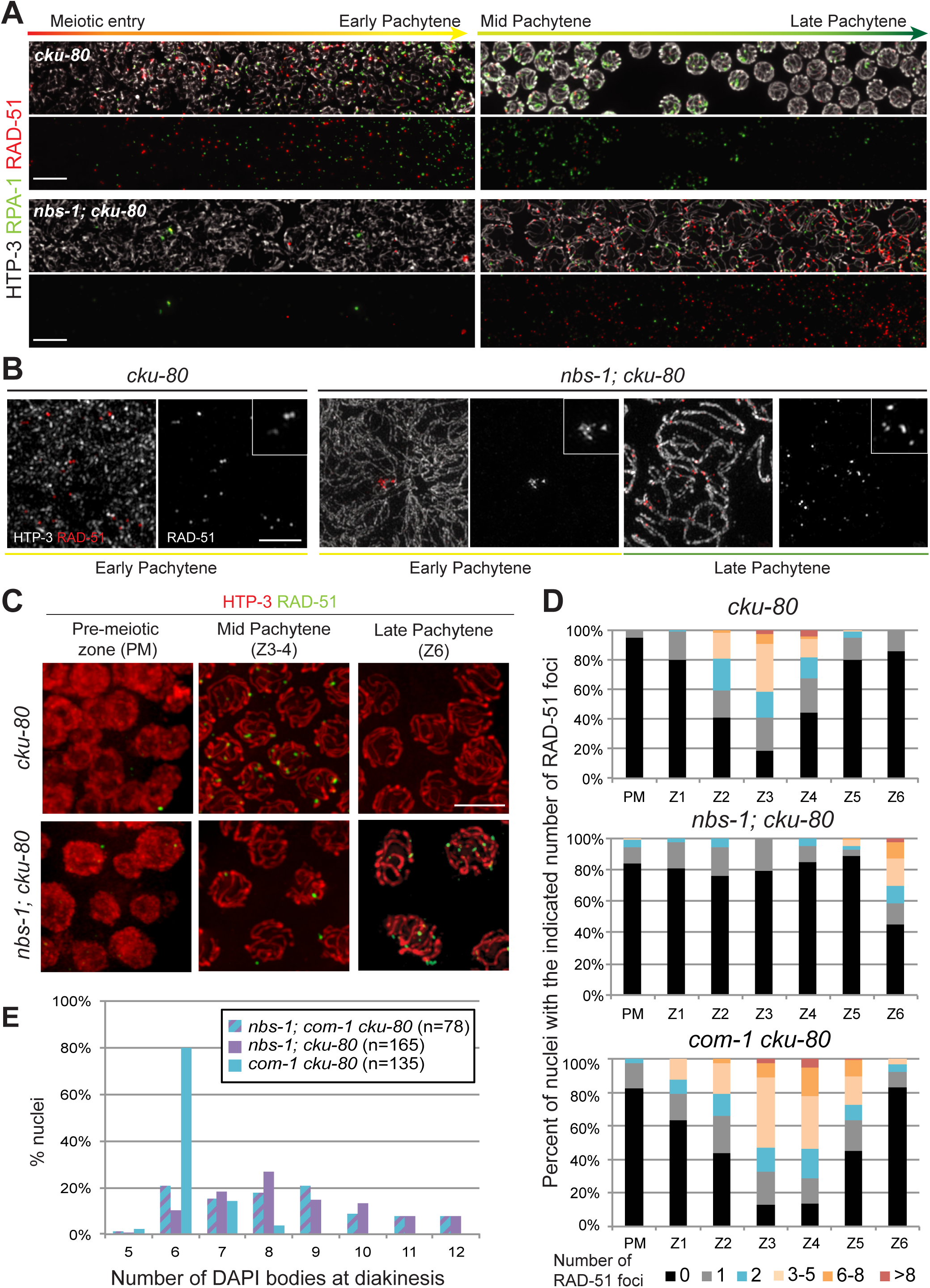
NBS-1 is required for timely loading of RAD-51 and RPA-1 during pachytene. **A)** Images of nuclei from spread gonads from *cku-80* and *nbs-1(me103); cku-80* mutant worms, immunostained for HTP-3 (greyscale), RAD-51 (red) and RPA-1 (green). Scale bar: 15 μm. The dynamics of appearance and removal of RAD-51 and RPA-1 in the *cku-*80 gonad are comparable to wild-type. In the *nbs-1; cku-80* double mutant RAD-51 and RPA-1 foci do eventually increase in abundance, in contrast to the *nbs-1* single mutant (Figure 3), but this rise only occurs in late pachytene and not in early pachytene as in the *cku-80* mutant or wild-type. **B)** SIM images of RAD-51 foci from the spread nuclei in (A). Inset image for *cku-80* shows the doublet (or singlet) configuration of RAD-51 foci (left), similar to wild type. In the *nbs-1; cku-80* double mutant, the RAD-51 foci present in early stages usually have a complex structure (center) but can appear as doublets (or singlets) in late pachytene (right). Scale bar: 2μm. **C)** Immunolocalization of HTP-3 (red) and RAD-51 (green) on whole-mount gonads Scale bar: 5μm. **D)** Quantification of the numbers of RAD-51 foci in whole-mount gonads, illustrating the contrast between the *nbs-1; cku-80* double mutant, in which increased abundance of RAD-51 foci is restricted to late pachytene (Zone 6) and the *com-1 cku-80* double mutant, which exhibits normal RAD-51 foci dynamics. **E)** Quantification of DAPI bodies in diakinesis nuclei, showing that successful bivalent formation occurs much more frequently in the *com-1 cku-80* double mutant than in either *nbs-1; cku-80* or *nbs-1; com-1 cku-80*.

This late timing of appearance of RAD-51 foci may help to explain why restoration of CO formation is incomplete in the *nbs-1; cku-80* double mutant. Initial loading of pro-CO factors must occur prior to the transition to late pachytene in order for DSB repair intermediates to become competent to mature into COs (19). Thus when resection is delayed, it may sometimes occur too late to enable recruitment of factors needed to generate COs.

### NBS-1 and COM-1 play distinct roles in promoting HR

Both COM-1 and NBS-1 are required for meiotic DSB repair but dispensable for DSB formation (Figure 2 and (13, 18)). However, our data indicate that their respective roles in resection and promotion of HR are quite different. Whereas elimination of *cku-80* resulted in a modest partial rescue of bivalent formation in the *nbs-1* background, with 10% of diakinesis nuclei showing 6 bivalents, we observed that loss of *cku-80* in the *com-1* background resulted in much more substantial restoration of bivalent formation, with 80% of diakinesis nuclei showing 6 bivalents (Figure 5E), recapitulating previous observations (13). Moreover, analysis of diakinesis nuclei in the *nbs-1; com-1 cku-80* triple mutant indicated that NBS-1 is required for the efficient bivalent formation observed in the *com-1 cku-80* mutant (Figure 5E). Together these results suggest that while COM-1 is required to antagonize CKU-80 and prevent NHEJ-mediated repair of DSBs, it is not essential for MRN-dependent resection to yield interhomolog COs.

This conclusion is further supported by comparison of RAD-51 dynamics in the *com-1 cku-80* and *nbs-1; cku-80* double mutants. In contrast to *nbs-1; cku-80* where RAD-51 foci did not increase in abundance until late pachytene, the *com-1 cku-80* double mutant exhibited RAD-51 foci dynamics similar to wild type, with a strong peak in foci numbers in mid pachytene and a decline in foci numbers by late pachytene (Figure 5D), as previously described (13). This indicates that COM-1 function is essential for resection in presence of NHEJ but becomes dispensable in the absence of NHEJ. This result implies that COM-1 is primarily required during meiosis to antagonize CKU-80 and NHEJ, but is not essential for timely MRN-dependent resection when NHEJ is abrogated (see also Discussion). In contrast, NBS-1 is required both for antagonizing NHEJ and for promoting resection.

### EXO-1 is required for CO formation and genome integrity, but not for late prophase RAD-51 loading in the *nbs-1; cku-80* double mutant

The presence of COSA-1 foci as well as late RAD-51 and RPA-1 foci in the *nbs-1; cku-80* double mutant made us wonder what factors might be mediating resection in this context. One candidate is the exonuclease Exo1, which has been shown to be involved alongside the MRN complex in promoting extended resection (9). Although EXO-1 is dispensable for meiotic recombination in otherwise wild-type *C. elegans* (13), EXO-1 is required for partial restoration of RAD-51 loading, CO formation and chromosome integrity in the *mre-11(iow1); cku-80* double mutant (14), indicating that delayed DSB resection and repair *via* HR are dependent on EXO-1 in this context. We therefore tested whether EXO-1 could mediate resection during late pachytene in the absence of NBS-1 (Figure 6).

**Figure 6:**
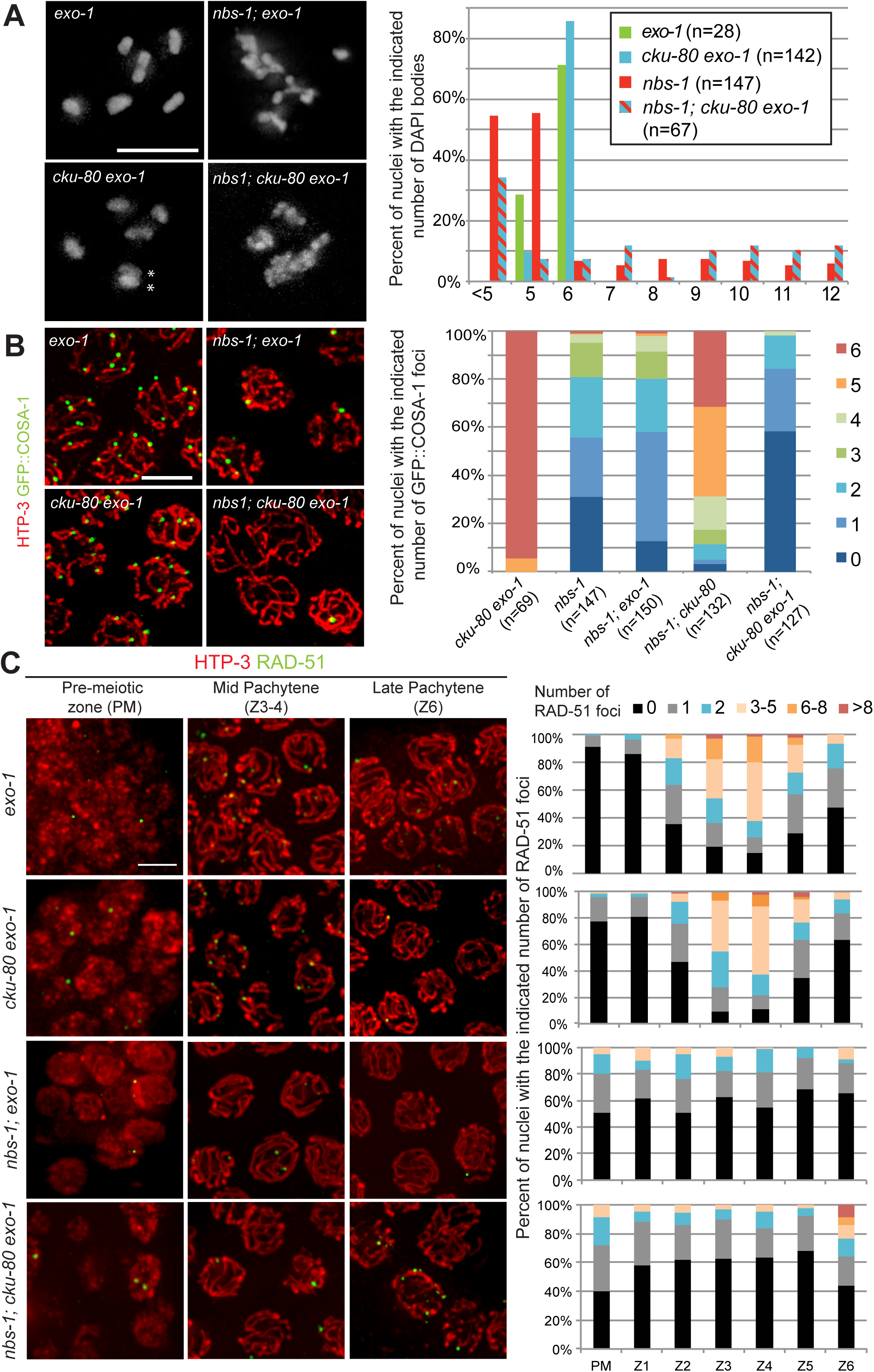
EXO-1 is required for DSB repair in the absence of cKU-80 and NBS-1. **A)** Left: Images of individual DAPI stained diakinesis nuclei (scale bar 5µm) showing the presence of chromosome aggregates in the *nbs-1; cku-80 exo-1* triple mutant. Asterisks in *cku-80 exo-1* indicate two chromosomes on top of each other. Right: Quantification of DAPI bodies in diakinesis nuclei. **B)** Left: Immunolocalization of HTP-3 (red) and COSA-1 (green) in late pachytene nuclei from whole-mount gonads. Right: Stacked bar graph representing the percentage of nuclei with the indicated numbers of COSA-1 foci in the different genotypes, showing that COSA-1 focus formation is eliminated in the *nbs-1(me102); cku-80 exo-1* triple mutant. **C)** Left: Immunolocalization of HTP-3 (red) and RAD-51 (green) in whole-mount preparations. Right: quantification of RAD-51 foci for the genotypes depicted on the left.

While the *nbs-1; cku-80* double mutant displayed mostly univalents and bivalents in diakinesis oocytes, we frequently observed chromosome aggregates at diakinesis in the *nbs-1; cku-80 exo-1* triple mutant (Figure 6A), suggesting partial redundancy of NBS-1 and EXO-1 function in maintaining genome integrity. Moreover, the partial rescue of GFP::COSA-1 focus formation observed in *nbs-1; cku-80* was also dependent on EXO-1, as the *nbs-1; cku-80 exo-1* triple mutant failed to form GFP::COSA-1 foci (Figure 6B). These results indicate a strict requirement for EXO-1 to form COs in the absence of both NBS-1 and NHEJ. However, EXO-1 was not essential for the late pachytene rise in RAD-51 foci observed in *nbs-1; cku-80* (Figure 6C), as a significant portion of late pachtyene nuclei (Z6) with numerous RAD-51 were detected in the *nbs-1; cku-80 exo-1* triple mutant. Persistence of late prophase RAD-51 foci but loss of CO site markers is reminiscent of phenotypes observed in the *com-1 cku-80 exo-1* triple mutant (13) and suggests either that the resection occurring in these contexts occurs too late for recruitment of CO factors or that EXO-1 has an additional late function in promoting CO formation, as has been observed in mouse and yeast (30, 31).

## Discussion

### Identification of *C. elegans* NBS-1 as a compact ortholog of NBS1/Xrs2

The MRN complex has long been recognized as a central player in mediating HR-based repair of DSBs across species, but there are substantial differences in the degree of conservation among its subunits (32). MRE11 and RAD50 are ancient in origin, are highly conserved among eukaryotes and have clearly identifiable orthologs in both eubacteria (SbcC and SbcD) and archae. In contrast, NBS1 orthologs are detected only in eukarya and are notoriously poorly conserved. Primary sequence conservation among orthologs from different kingdoms is mainly restricted to the N-terminal FHA domain, and conservation outside this domain is marginal even within kingdoms, *e.g. S. cerevisiae* Xrs2 and *S. pombe* Nbs1 share only 10% identity in the 250 amino acids following the FHA domain, and the presence of tandem BRCT domains within this region had remained unrecognized in many orthologs until introduction of an algorithm specifically designed to detect such motifs (15). Indeed, when human NBS1 was first discovered, its protein size and association with MRE11 and RAD50 were crucial for recognizing NBS1 and Xrs2 as functional homologs (33).

Although *C. elegans* MRE-11 and RAD-50, and their roles in meiotic recombination and DNA repair, have been known for some time (14, 16, 17), the nematode counterpart of NBS1/Xrs2 had remained elusive. Our identification of C09H10.10 as *C. elegans* NBS-1 makes the reason it had escaped detection apparent: while it contains both the N-terminal FHA domain and the conserved MRE-11 interaction domain (MID) near its C-terminus, *Ce* NBS-1 is only about half the size of most other NBS1 orthologs and lacks the tandem BRCT domains. This stripped-down version of NBS-1 present in *C. elegans* is nonetheless sufficient to support the functions of MRE-11 and RAD-50 in promoting efficient and timely meiotic DSB repair and in repairing/preventing accumulation of replication-associated DNA damage. The fact that such a compact version of NBS1 can support the essential functions of MRE-11 and RAD-50 in DSB repair parallels the recent finding that a 108 amino acid fragment of mammalian Nbs1 (which contains the MID but lacks both the tandem BRCT motifs and the N-terminal FHA domain) can substantially support essential functions of Mre11 and Rad50 in mouse cells *in vitro* and *in vivo* (34).

### NBS-1-independent functions of MRE11 and RAD50 during *C. elegans* meiosis

In all species where it has been studied, the MRN complex has been shown to be crucial for repair of meiotic DSBs (35). However, involvement of MRN in the formation of such breaks varies from species to species. Whereas Mre11, Rad50 and Nbs1 are not required for meiotic DSB formation in *S. pombe* or *A. thaliana* (36–40), all three core members of MRX are required for DSB formation during *S. cerevisiae* meiosis (41, 42). Interestingly, our analysis here revealed that these two meiotic functions of MRN complex components can be uncoupled. While *C. elegans* NBS-1 is integral to the functions of the MRN complex in promoting timely resection and repair of meiotic DSBs, we found that the previously reported roles of MRE-11 and RAD-50 in promoting DSB formation (16, 17) do not require NBS-1 (Figure 7).

**Figure 7:**
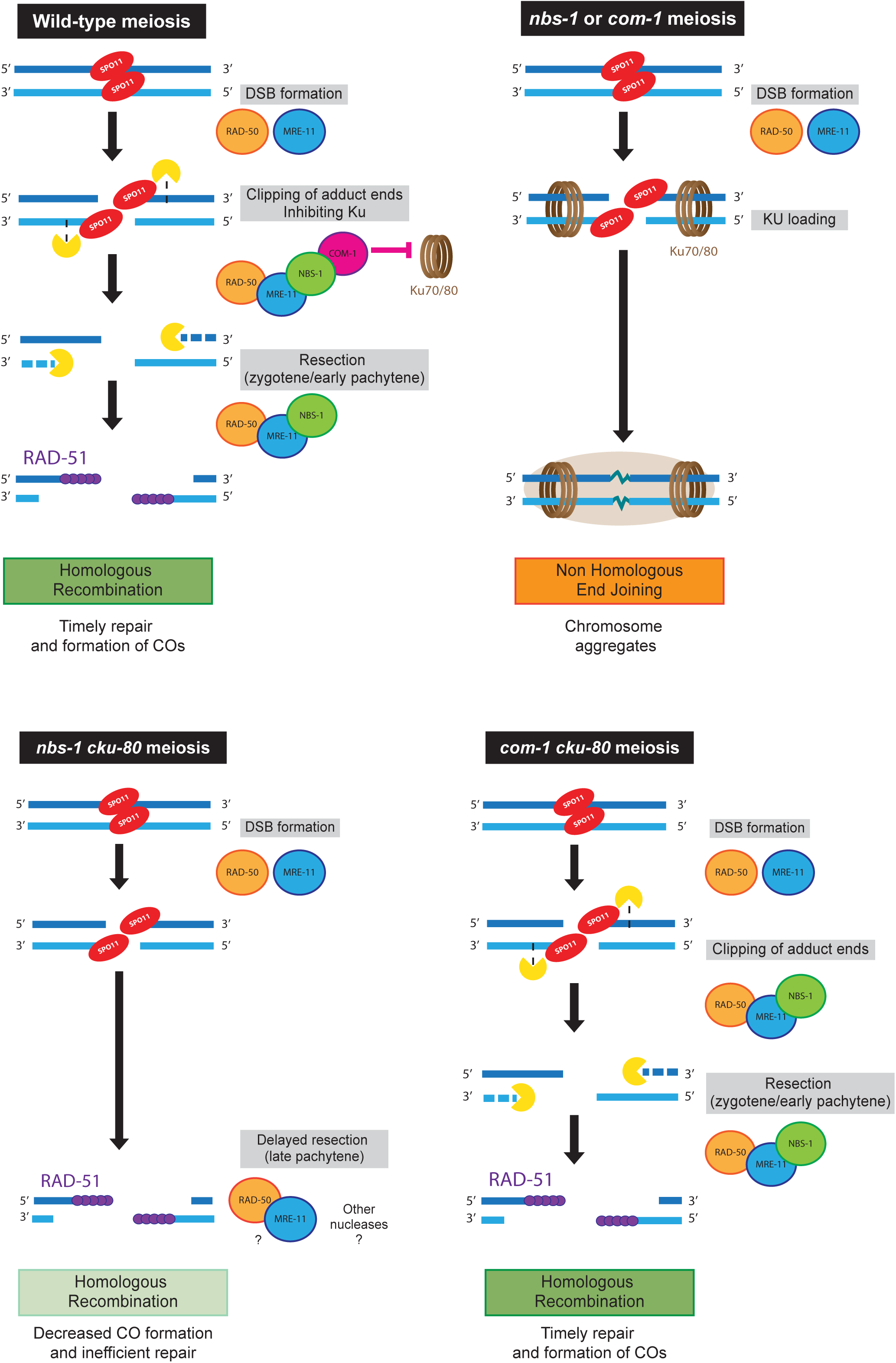
NBS-1 is essential for COM-1 recruitment to inhibit NEHJ and for MRN function to promote timely resection of DSBs at meiosis. Working model derived from the findings of this study and others, as discussed in the text.

How MRN complex components function to promote DSB formation remains unknown. However, separation-of-function mutations that uncouple DSB formation and repair activities may be informative. Missense mutations in *C. elegans* [*mre-11(iow1*)] and *S. cerevisiae* [*mre11-D16A*] that impair DSB resection but not DSB formation affect the same conserved phosphoesterase domain and destabilize the interaction between MRE11 and RAD50 ((14, 43, 44) and this study). This suggests that a stable interface between MRE11 and RAD50 that is essential for resection and repair activities of the MRN complex may be less important for DSB-promoting activity, raising the possibility that MRE11 and RAD50 may function in a different conformation (34, 45) or stoichiometry (46), or even as separate proteins, to influence DSB formation. Further, in *S. cerevisiae*, Xrs2 may be required for DSB formation partially based on its role in promoting nuclear localization of Mre11 (47); conversely, the fact that *C. elegans* NBS-1 is dispensable for DSB formation indicates that (at least some) MRE-11 and RAD-50 must get into the nucleus without NBS-1.

Additional evidence suggests that *C. elegans* MRE-11 may also be able to function independently of NBS-1 in another context. Specifically, we found that late RAD-51 foci reflecting delayed end resection were present in late pachytene nuclei in *nbs-1; cku-80 exo-1* mutant germ lines, whereas such foci were absent in *mre-11; cku-80 exo-1* (14). This result suggests that MRE-11 may be capable of promoting some degree of end resection in late pachytene nuclei in the absence of NBS-1 and EXO-1.

### Distinct roles for MRN and COM-1 in promoting DSB resection and antagonizing NHEJ

DSBs pose a threat to genome integrity, and DNA repair machineries have evolved to prevent or limit their damaging consequences. Moreover, evidence for competition between different DSBR pathways is present in all studied species. For example, elimination of Ku in mammalian cells increases the frequency of DSB-induced HR between direct repeats (48), and conversely, elimination of Mre11 results in higher incidence of NHEJ in yeast cells (49). The extent to which DSBs are repaired using mutagenic repair mechanisms such as NHEJ *vs.* high-fidelity mechanisms such as HR depends on cellular context. During meiosis, it is crucial that DSBs be repaired strictly by HR, both (i) to promote the formation of interhomolog COs needed to segregate chromosomes and (ii) to restore genome integrity while minimizing introduction of new mutations. However, even during meiosis where the outcome of DSB repair is so heavily biased toward HR, abrogation of HR in *C. elegans* germ cells has revealed that NHEJ factors are nevertheless still present and can promote illegitimate repair ((13, 14) and this study). The MRN complex and COM-1 are crucial during meiosis to tip the balance irrevocably toward the HR outcome.

The current work, integrated with previous findings (13, 14), demonstrates that *C. elegans* MRN and COM-1 make distinct contributions to promoting HR and antagonizing NHEJ during meiosis (Figure 7). NBS-1, MRE-11 and COM-1 are all required to prevent meiotic catastrophe resulting from inappropriate engagement of NHEJ. However, in the absence of Ku, differences in the roles of these components are revealed. When Ku is removed in an *nbs-1* or *mre-11(iow1)* mutant background, RAD-51 loading (indicative of end resection) is delayed and CO formation is inefficient, However, when Ku is removed in a *com-1* mutant background, RAD-51 foci levels and timing appear normal and CO formation is much more efficient. These findings indicate that COM-1 is required primarily to antagonize Ku, yet is substantially dispensable for MRN-mediated end resection when Ku is absent (Figure 7). Whereas MRN can promote efficient and timely end resection without COM-1 (in combination with EXO-1; see below), however, MRN cannot function without COM-1 to antagonize Ku. We interpret these findings in light of reports that the *S. pombe* Nbs1 FHA domain directly engages Ctp1 and that Ctp1/CtIP is recruited to DSB sites through NBS1 in both *S. pombe* and human cells (50–52). Specifically, we propose that during *C. elegans* meiosis, NBS-1 couples resection initiation and inhibition of NHEJ both by participating in MRN-mediated end-resection and by recruiting COM-1 to DSB sites. Further, based on structural analysis of *S. pombe* Ctp1 suggesting an ability to form bridges between MRN-C complexes on opposite sides of a DSB (53, 54), we propose that MRN-C may play a dual role in antagonizing NHEJ both by promoting endonucleolytic cleavage to initiate resection and by mediating bridging between DNA ends, thereby preventing the loading of the pre-formed Ku ring.

### Redundancy in HR machinery contributes to robustness of repair

Genome integrity of germ cells is paramount to perpetuation of species. As faithful chromosome inheritance during sexual reproduction depends on meiotically-induced DSBs, it is crucial that DSB repair in germ cells be highly robust. Synthesis of the current work with prior analyses of MRN-C function in the *C. elegans* germ line suggests that partial redundancy among factors and activities promoting DSB resection may contribute to robustness of the system.

From the onset of meiotic prophase through the end of the early pachytene stage, DSB end resection is highly dependent on MRN (14, 17) and this study). As DSBs must be processed and engage the homolog during early prophase in order to be competent for CO formation (19), the timely participation of MRN in DSB resection is thus crucial for efficient CO formation. In contrast, *C. elegans* EXO-1 is not required for meiotic DSB resection in otherwise wild-type germ cells. However, EXO-1 can mediate resection during late prophase in the absence of MRN activity, and either MRN or EXO-1 can mediate resection during late prophase in the absence of Ku. Further, in a *com-1 cku-80* double mutant, DSBs can undergo timely resection during early prophase, but now both MRN and EXO-1 are required for this to occur. This indicates that EXO-1 is available during early prophase and can augment resection either through its own exonuclease function or by enhancing MRN activity when the system is compromised by loss of COM-1. We suggest that although EXO-1 is largely dispensable for successful meiosis, it likely does collaborate with MRN-C during normal meiosis to help ensure a reliable outcome.

## Material and Methods

### Strains and genetics

All C. elegans strains were cultivated at 20°C under standard conditions. Strains used in this study are:

AV630 *meIs8 [gfp::cosa-1] II*

AV727 *meIs8 [gfp::cosa-1] II*, *ruIs32* [*unc-119*(+); *pie-1::mcherry::histoneH2B*] *III*; *itIs38*

[*pAA1; pie-1::GFP::PH::unc-119(+)*]

AV828 *nbs-1(me102) meIs8/mIn1* [*mIs14 dpy-10(e128)*] *II*; backcrossed 3x from the original balanced strain

AV845 *spo-11(me44)/ nT1 [unc(n754dm) let] IV*

AV846 *nbs-1(me102) meIs8/mIn1* [*mIs14 dpy-10(e128)*] *II; spo-11(me44)/ nT1 IV*

AV860 *nbs-1(me103)/mIn1* [*mIs14 dpy-10(e128)*] *II*

AV861 *nbs-1(me104)/mIn1* [*mIs14 dpy-10(e128)*] *II*

AV862 *nbs-1(me105)/mIn1* [*mIs14 dpy-10(e128)*] *II*

AV863 *nbs-1(me106)/mIn1* [*mIs14 dpy-10(e128)*] *II*

AV865 *nbs-1(me102) meIs8/mIn1* [*mIs14 dpy-10(e128)*] *II; mre-11(ok179)/ nT1 V*

AV874 *meIs8 II; cku-80(ok861) III*

AV875 *meIs8 II; exo-1(tm1842) III*

AV876 *meIs8 II; cku-80(ok861) exo-1(tm1842) III*

AV877 *nbs-1(me102) meIs8/mIn1*[*mIs14 dpy-10(e128)*] *II; cku-80(ok861) III*

AV878 *nbs-1(me102) meIs8/mIn1* [*mIs14 dpy-10(e128)*] *II; exo-1(tm1842) III*

AV879 *nbs-1(me102) meIs8/mIn1* [*mIs14 dpy-10(e128)*] *II; cku-80(ok861) exo-1(tm1842) III*

AV904 *nbs-1(me103)/mIn1* [*mIs14 dpy-10(e128)*] *II; opIs263[rpa-1::yfp, unc-119+]*

AV905 *cku-80(ok861) III; opIs263[rpa-1::yfp, unc-119+]*

AV947 *nbs-1(me103)/mIn1* [*mIs14 dpy-10(e128)*] *II; cku-80(ok861) III; opIs263[rpa-1::yfp, unc-119+]*

XF0644 *com-1(t1626) unc-32(e189)/hT2 [bli-4(e937) let-?(q782) qIs48] III*

XF0697 *com-1(t1626) unc-32(e189 cku-80(tm1524) / hT2 cku-80(tm1524) III*

### nbs-1(me102) isolation, mapping and identification

*nbs-1(me102)* was isolated in a genetic screen for meiotic mutants exhibiting altered numbers and/or appearance of GFP::COSA-1 foci. The AV727 strain used for this screen allowed simultaneous live imaging of GFP::COSA-1 foci, chromatin (mCherry::H2B) and germ cell membranes (GFP::PH). F1 progeny of EMS mutagenized parents were plated individually, and pools of adult F2 progeny from each F1 plate were mounted on multi-well slides in anesthetic (0.1% tricaine and 0.01% tetramisole in M9 buffer) to visualize their germ lines; candidate mutations were recovered from siblings of visualized worms. The *me102* mutation was balanced by the *mIn1 II* balancer, then mapped to a ~6.8cM region on chromosome *II* between *unc-4* and *rol-1*. Following backcrossing (3x) to generate the AV828 strain, homozygous *me102* worms were subjected to whole genome sequencing. DNA was extracted from approximately 400 individually picked *me102* homozygous or AV727 gravid adult worms, which were rinsed twice in M9 and resuspended in 10mM EDTA, 0.1M NaCl. Worms were then pelleted, flash frozen in liquid nitrogen and resuspended in 450μL of lysis buffer containing 0.1M TRIS pH 8.5, 0.1M NaCl, 50mM EDTA and 1% SDS plus 40 μL of 10mg/ml proteinase K in TE pH 7.4, vortexed, and incubated at 62°C for 45 minutes. Two successive phenol-chloroform extractions were performed using the Phase Lock gel tubes from Invitrogen, and DNA was precipitated with 1mL of 100% ethanol plus 40 μL of saturated NH_4_Ac (5M) and 1μL of 20 mg/ml GlycoBlue. The DNA pellet was washed with 70% ethanol, air-dried and resupended in 50μL of TE pH 7.4. Paired-end libraries were prepared using the Nextera technology (Illumina) and sequencing was performed on a MiSeq sequencer (2x75bp). Reads were mapped to *C. elegans* reference genome (WBcel 235) using Bowtie software. Variant calling was performed using UnifiedGenotyper software from GATK (https://software.broadinstitute.org/gatk) and lists from AV828 and AV727 were compared to eliminate non-causal SNPs and INDELS. Two mutations in the 6.8Mb interval on chromosome II were specific to the *me102* strain. Both were canonical EMS induced G>A or C>T mutations, one a missense mutation in the *C07E3.3* gene and the other a nonsense mutation in the *C09H10.10* gene.

### CRISPR genome editing

We used direct injection of Cas9 protein (PNAbio) complexed with sgRNA generated by *in vitro* transcription from a PCR template. crRNAs were designed using either Benchling (benchling.com) or ChopChop (chopchop.rc.fas.harvard.edu), following guidelines from (55). crRNAs used to generate *me103, me104, me105* and *me106* alleles were GAGCATAGAATGGGGCGATG and GTTCATGCGAGCATAGAATG (see also Figure S1). dsDNA template for RNA transcription was obtained by PCR amplification using a "universal" reverse primer (oCG83: AATTTCACAAAAAGCACCGACTCGGTGCCACTTTTTCAAGTTGATAACGGACTAGCCTTATTTTAACTTGCTATTTCTAGCTCTAAAAC) and a forward primer containing the T7 promoter sequence upstream of the crRNA sequence as well as 20bp of complementarity with oCG83 (namely oCG84 TAATACGACTCACTATAGGG-GAGCATAGAATGGGGCGATGGTTTTAGAGCTAGAAAT; and oCG85:

TAATACGACTCACTATAGGGGTTCATGCGAGCATAGAATGGTTTTAGAGCTAGAAAT). PCR was performed with the Phusion master mix from NEB in 50 μL with 4 μL of each oligo (10mM stock), using the following program: 94°C for 5 min; then 25 cycles of: 94°C for 30 seconds, 55°C for 30 seconds, 72°C for 30 seconds; ending by a step at 72°C for 5 mins. dsDNA was purified on column, and concentration assessed by Nanodrop. In vitro transcription was done overnight using the Ambion MEGAscript Kit from ThermoFisher. Ensuing RNA purification was performed using the MEGAclear Kit with a final elution volume of 40 μL. Cas9/sgRNA complexes were formed for 10min at room temperature with 500ng/μL of Cas9 protein (PNABio) and 250ng/ μL of both sgRNA (total final concentration for both guides combined). N2 worms (P0) were injected with the mix along with pCJF104 as a co-injection marker (56). Red F1s (carrying pCJF204) were singled out, and a subset of F2 progeny were fixed and stained with DAPI (see below) to assess the phenotype of diakinesis nuclei. From plates containing worms exhibiting aggregated chromosomes at diakinesis, the new mutations were recovered from siblings of the imaged worms and balanced by *mIn1*. The *nbs-1* locus was amplified from homozygous mutant worms using oCG48 (GAGAAAGGCTCCGTGGTCAA) and oCG50 (GCCGTCAACTTCCAGAGTCA) primers and subjected to Sanger sequencing (Sequetech, 935 Sierra Vista Ave. Ste. C, Mountain View, CA 94043). Details of the mutations can be found in Figure S1.

### Yeast two-hybrid experiments

Worm RNAs were extracted by adding 250µL Trisol to 20µL wild-type N2 worm pellet in M9 and incubation at 4°C for 30min, followed by standard phenol-chloroform extraction (see above). cDNAs were obtained from these RNAs using the Superscript III First-strand synthesis for RT-PCR by Invitrogen. cDNA sequences of MRE-11, RAD-50, COM-1 and NBS-1 were amplified using the following primers containing SpeI (blue) and AvrII (red) restriction sites to allow for cloning into pDP133 (prey vector, complementing leucine auxotrophy) and pDP134-135 (bait vectors, complementing tryptophan auxotrophy) (57) with following primers:

MRE11 forward: NNNNACTAGTATGTGTGGCAGTGA

MRE11 reverse: NNNNCCTAGGTTAGAAGAAACTTAG

RAD-50 forward: CTAACTAGTATGGCGAAATTTTTACGCCTACAC

RAD-50 reverse: CTACCTAGGGAACCGTCTCTTCGTATTAACTCT COM-1 forward: NNNNACTAGTATGCAATCTGTGGATCCATTTG

COM-1 reverse: NNNNCCTAGGTTAATTCCACGTATTGATTCCAGTCGG

NBS-1 forward: NNNNACTAGTATGCCCATCAATGGCATAAAAATCAAAAACTC

NBS-1 reverse: NNNNCCTAGGTCAGTGCACAATTCT.

The plasmid bearing the mutated version of MRE-11 (MRE-11 *iow1*) was generated using Gibson assembly (NEB) to replace a 366 bp SpeI XbaI fragment from pDP133-MRE11 with a corresponding dsDNA fragment containing the iow1 mutation.

Yeast strain YCK580 was transformed according to (58) with plasmid pairs with one plasmid containing the prey fused with the GAL4 activation; the other containing the bait fused with the LexA DNA binding domain. Transformed cells were spread on selective media lacking both leucine and triptophan (-LW) and grown for 48h. One clone was selected for each pair, and interaction was assayed on media lacking histidine (- LWH) with or without the His3p competitive inhibitor 3-AT (25mM).

### Cytological analysis

Numbers of DNA bodies present in diakinesis oocytes were assessed in intact adult hermaphrodites at 24h post L4 stage, fixed in ethanol and stained with 49,6-diamidino-2-phenylindole (DAPI) as in (59). Immunostaining for GFP::COSA-1 and RAD-51 in whole-mount gonads was conducted as in (60). All experiments were performed on gonads dissected at 24-26h hours post L4 at 20°C. The following primary antibodies were used at the indicated dilutions in PBS with 0.1% Tween: chicken anti-HTP-3 (1:500 (61)); rabbit anti-GFP (1:200 (19)), rat anti-RAD-51 (1:500 (62)). Dual RPA-1::YFP/RAD-51 immunostaining was performed on spread gonads as in (63), with YFP being detected by the rabbit anti-GFP antibody. All images were acquired using a 100x NA 1.40 objective on a DeltaVison OMX Blaze microscopy system, deconvolved and corrected for registration using SoftWoRx. Gonads were subsequently assembled using the “Grid/Collection” plugin (64) in ImageJ. Wide field images were obtained as 200 nm spaced Z-stacks, while 3D-SIM images were obtained as 125 nm spaced Z-stacks. For display, contrast and brightness were adjusted in individual color channels using ImageJ. For quantification of RAD-51 foci, at least 3 gonads were counted per genotype. Gonads were divided into 7 zones: the premeiotic zone (PM), where HTP-3 appears diffuse in the nulei, and into 6 equal-sized zones based on physical distance from meiotic entry (where HTP-3 signal forms tracks along chromosome length) to late pachytene (end of cell rows). For the GFP:COSA-1 experiments, nuclei within the last 6 cell rows were counted; numbers of nuclei counted were as follow: wild type (n=115), *cku-80* (n=145), *exo-1* (n=205), *cku-80 exo-1* (n=69), *nbs-1(me102)* (n=147), *nbs-1; cku-80* (n=132), *nbs-1; exo-1* (n=150), *nbs-1; cku-80 exo-1* (n=127).

### Gamma-irradiation

Worms were exposed to 5kRad (50Gy) of γ-irradiation using a Cs-137 source at 20 hours post L4 stage. RAD-51 immunostaining was performed on gonads dissected and fixed 1h after irradiation, diakinesis DAPI body counts were done using worms fixed at 18h to 20h post irradiation.

## Acknowledgments

We thank Nimit Jain for help with sequence analysis, Stephanie Tzouanas & Stephanie Zimmerman for help with CRISPR methods, and Alex Woglar for instruction regarding chromosome spreading and SIM imaging, Chantal Akerib, Sree Ramakrishnan and Alex Woglar for comments on the manuscript. We thank S. Smolikove, V. Jantsch, A. Dernburg, M.Tijsterman and the Caenorhabditis Genetics Center (funded by NIH Office of Research Infrastructure Programs P40 OD010440) for reagents and strains and Raphael Guerois for helpful discussions. We also thank the Stanford Cell Sciences Imaging Facility, which is partially funded by Award Number 1S10OD01227601 from the National Center for Research Resources (NCRR)

This work was supported by a CJS fellowship from INRA and an award from the Bettencourt-Schueller Foundation to CG and by NIH grant R01GM67268 and American Cancer Society Research Professor Award RP-15-209-01-DDC to AMV. The contents of this manuscript are solely the responsibility of the authors and do not necessarily represent the official views of the funders, who had no role in study design, data collection and analysis, decision to publish, or preparation of the manuscript.

---

Supplemental information

for Girard *et al.* "Interdependent and separable functions of *C. elegans* MRN-C complex members couple formation and repair of meiotic DSBs"

**Figure S1:**
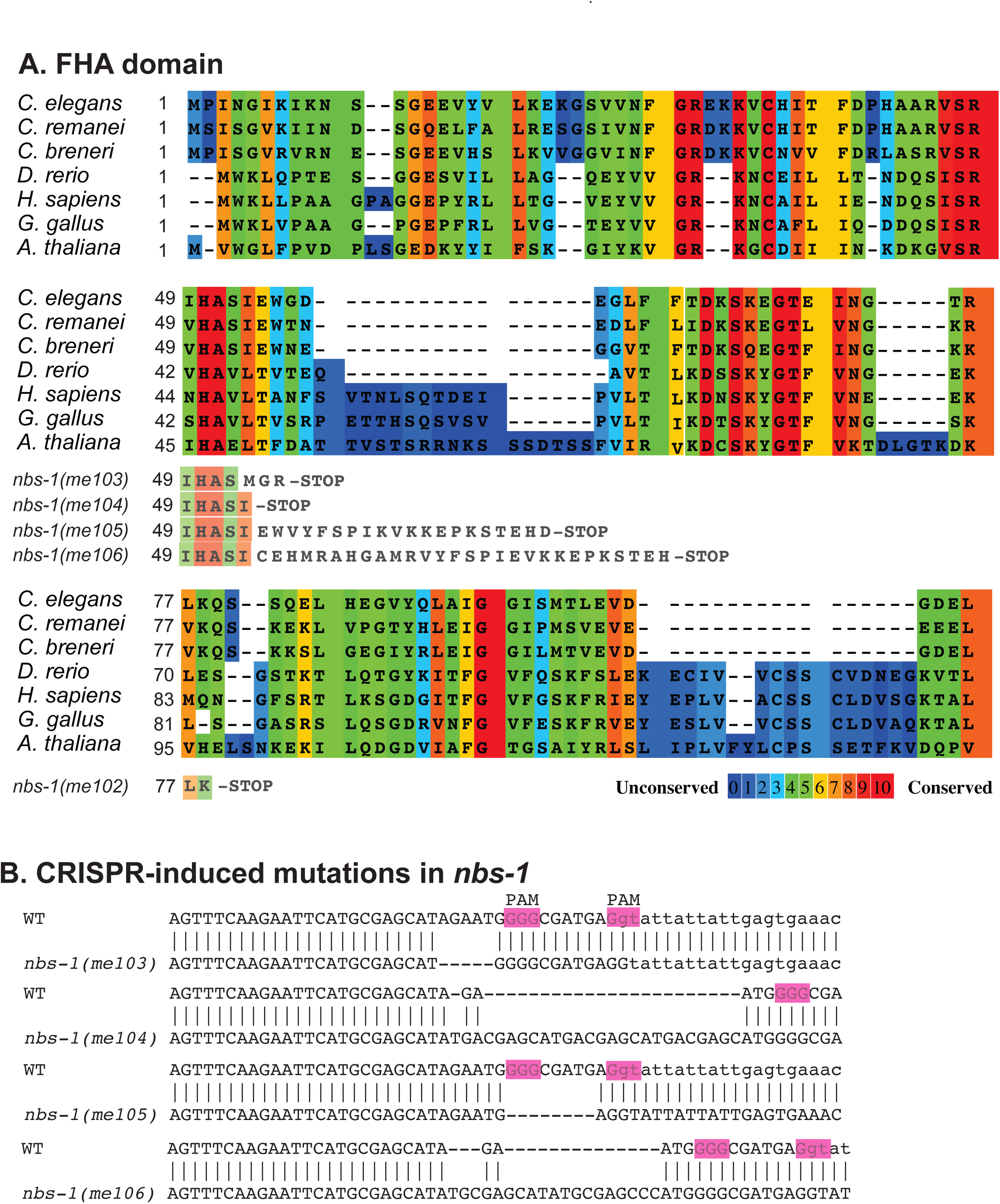
A) T-Coffee alignment showing conservation of the FHA domain among members of the NBS1/Nibrin protein family, with color coding generated using the PRALINE software. Predicted effects of *nbs-1* mutations obtained through EMS mutagenesis (*me102*) or CRISPR mutagenesis (*me103-me106*) on the protein sequence are also shown. Accession numbers are: *H. sapiens* BAA28616.1*, G. gallus* NP_989668.1*, *D.* rerio* NP_001014819.1*, S. cerevisiae* AAA35220.1*, S. pombe* BAC80248.1*, C. elegans* NP_496374.2 **B)** Genomic DNA sequence alignments between the wild-type *nbs-1* gene (starting at 179bp after ATG) and *nbs-1* mutant alleles generated by CRISPR. The PAM sequences (NGG) targeted are indicated in pink.

**Figure S2:**
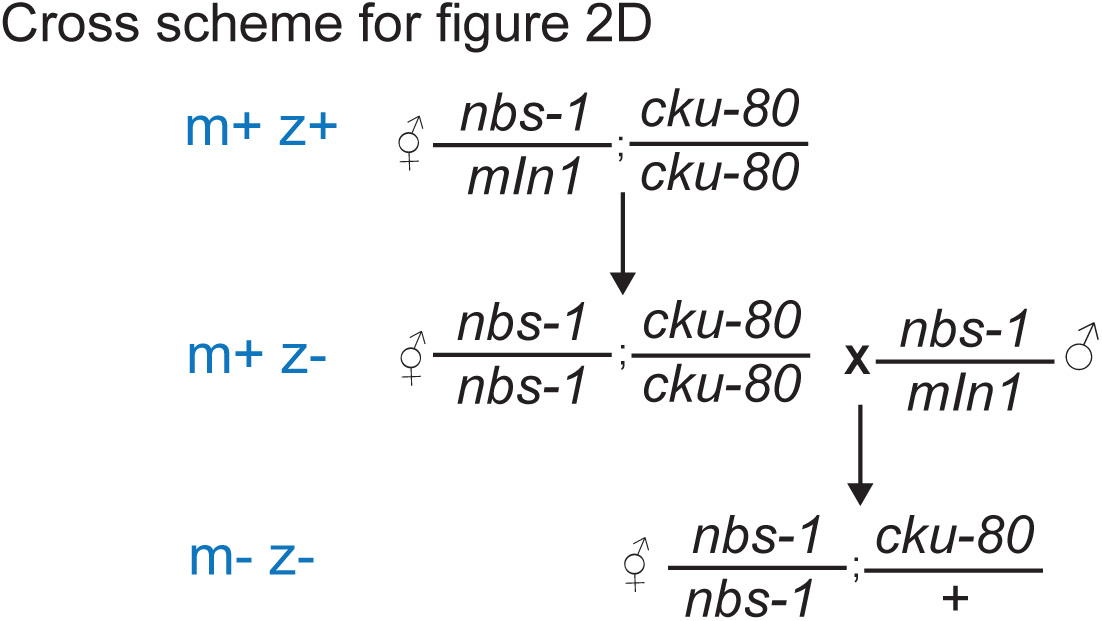
Crossing strategy used to obtain homozygous *nbs-1(me102)* mutant worms from homozygous *nbs-1(me102)* mothers (m-z-). *mIn1* refers to the balancer chromosome used to maintain the *nbs-1* mutation in a heterozygous state. Because progeny viability is partially restored in the *nbs-1; cku-80* double mutant (Table 1), viable m-z- *nbs-1* homozygotes that contained a wild-type *cku-80(+)* allele could be generated.

**Figure S3:**
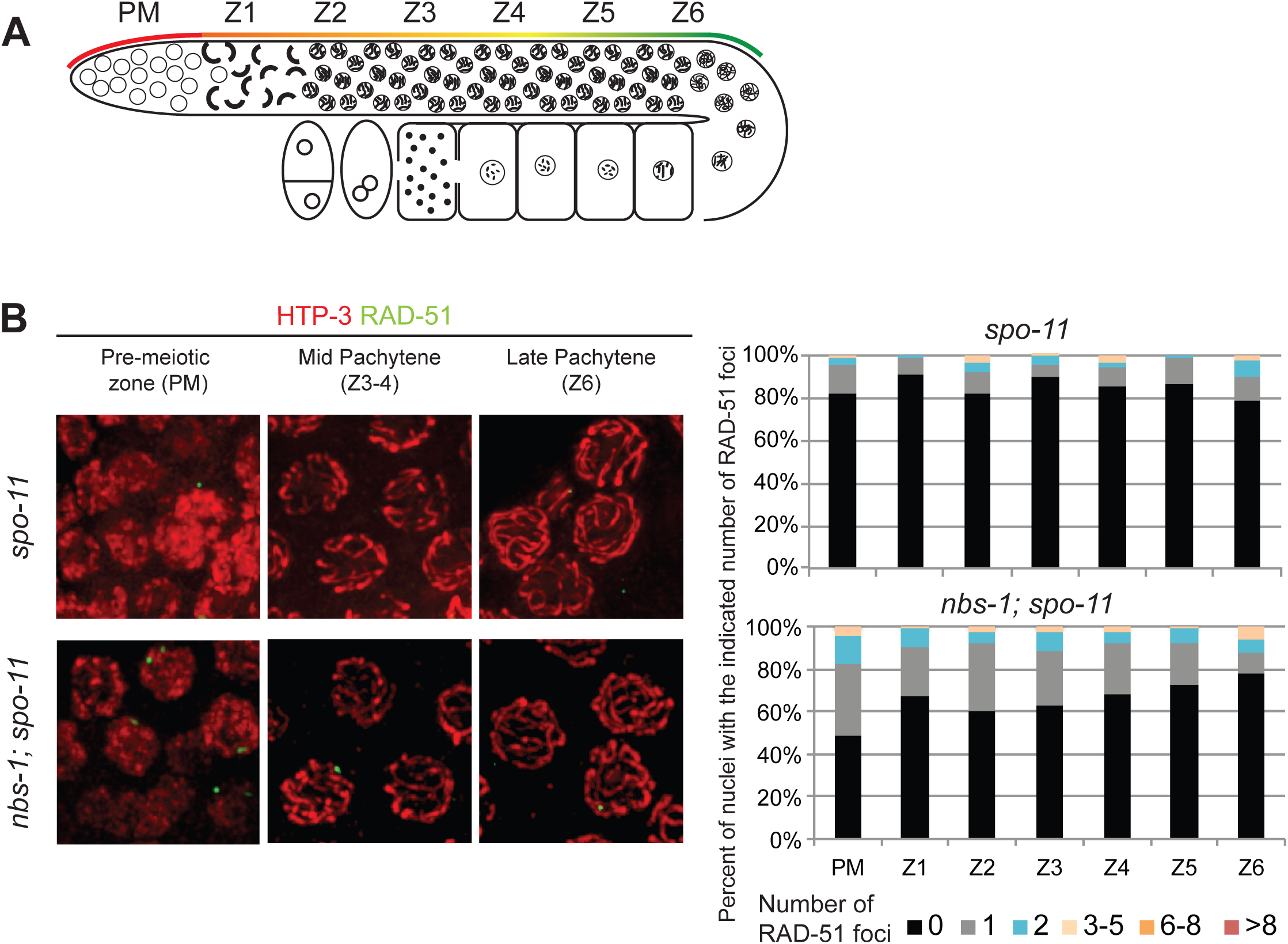
A) Schematic representation of the spatio-temporal organization of *C. elegans* gonad and the 7 consecutives zones from the pre-meiotic tip (PM) through the end of pachytene (Z6) used to assess RAD-51 foci numbers in the different mutants throughout this study (see Material and Methods for more details). **B)** Left: Immunolocalization of HTP-3 (red) and RAD-51 (green) in nuclei from whole-mount gonads. Right: Quantification of RAD-51 foci in the seven consecutive gonad zones defined in (A) in both *spo-11* and *nbs-1; spo-11* mutants.

**Figure S4:**
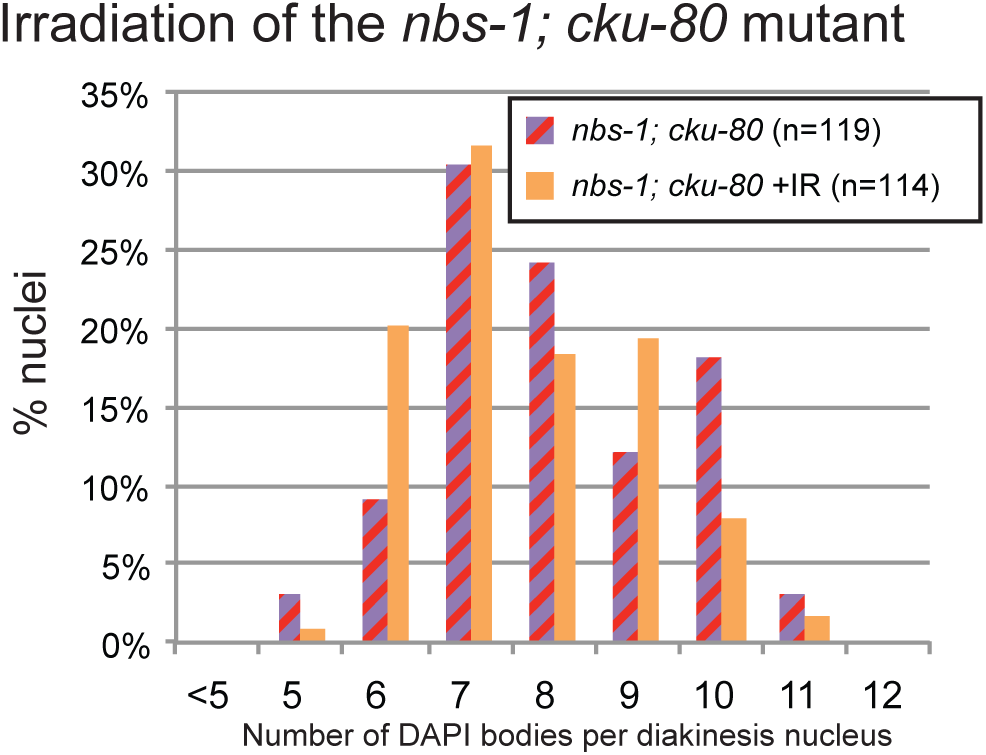
Quantification of DAPI bodies in diakinesis nuclei in the *nbs-1; cku-80* double mutant exposed to 5kRad γ-irradiation (mean 7.7 ± 1.3; n=114) and in the unirradiated control (7.9 ± 1.4, n=119, Mann-Whitney p=0.21). Partial restoration of CO formation in *nbs-1; cku-80* is not improved by excess DSB formation.

